# The genetic impact of an Ebola outbreak on a wild gorilla population

**DOI:** 10.1101/2021.05.31.446409

**Authors:** Claudia Fontsere, Peter Frandsen, Jessica Hernandez-Rodriguez, Jonas Niemann, Camilla Hjorth Scharff-Olsen, Dominique Vallet, Pascaline Le Gouar, Nelly Ménard, Arcadi Navarro, Hans R. Siegismund, Christina Hvilsom, M. Thomas P. Gilbert, Martin Kuhlwilm, David Hughes, Tomas Marques-Bonet

## Abstract

**Background:** Numerous Ebola virus outbreaks have occurred in Equatorial Africa over the past decades. Besides human fatalities, gorillas and chimpanzees have also succumbed to the fatal virus. The 2004 outbreak at the Odzala-Kokoua National Park (Republic of Congo) alone caused a severe decline in the resident western lowland gorilla (*Gorilla gorilla gorilla*) population, with a 95% mortality rate. Here, we explore the immediate genetic impact of the Ebola outbreak in the western lowland gorilla population.

**Results:** Associations with survivorship were evaluated by utilizing DNA obtained from fecal samples from 16 gorilla individuals declared missing after the outbreak (non-survivors) and 15 individuals observed before and after the epidemic (survivors). We used a target enrichment approach to capture the sequences of 123 genes previously associated with immunology and Ebola virus resistance and additionally analyzed the gut microbiome which could influence the survival after an infection. Our results indicate no changes in the population genetic diversity before and after the Ebola outbreak, and no significant differences in microbial community composition between survivors and non-survivors. However, and despite the low power for an association analysis, we do detect six nominally significant missense mutations in four genes that might be candidate variants associated with an increased chance of survival.

**Conclusion:** This study offers the first insight to the genetics of a wild great ape population before and after an Ebola outbreak using target capture experiments from fecal samples, and presents a list of candidate loci that may have facilitated their survival.

## Background

The Ebola virus (EBOV), discovered in 1976, causes a severe disease and often fatal hemorrhagic fever for which numerous human outbreaks have been reported throughout Africa [1]. The most virulent outbreak reported to date was in West Africa in December 2013 and lasted until 2016 with more than 28,000 confirmed or suspected human cases and more than 11,000 human deaths [2]. Since then, other outbreaks of Ebola have been observed. In June 2020, when the 2018 outbreak was declared over by the World Health Organization (WHO), 3,470 cases had been reported with 2,287 deaths (fatality rate of 66%) [3].

EBOV belongs to the single-stranded RNA virus family *Filoviridae* [4] with five distinct strains in the Ebola genus: *Zaire, Sudan, Budibugyo, Taï Forest* and *Reston*. The first three are responsible for the majority of human infections [5, 6]. The virus is highly infectious and can enter the body through direct contact of broken skin or mucous membranes with infected blood or body fluids, causing symptoms including fever, vomiting, diarrhea, internal and external bleeding. Ebola hemorrhagic fever or Ebola virus disease (EVD) is an acute and severe disease with a fatality rate in humans around 50% [5–7].

Infectious diseases such as Ebola are considered to be a threat to the survival of African great apes [8], together with other threats such as habitat loss, climate change and poaching [9]. In some cases, the human outbreaks have been linked to contact with infected bushmeat from gorillas or chimpanzees [10] and several surveys have reported dramatic declines in populations of great apes in parallel with human EVD outbreaks with laboratory confirmation of Ebola virus infection in some carcasses [10–12]. Gorilla populations from the Republic of Congo suffered severe die-offs during a chuman EVD outbreak near the Lossi sanctuary in 2002-2003 [12] and Odzala-Kokoua National Park in 2004 [13] with reported mortality rates as high as 95%. In Lossi sanctuary alone, it was estimated that the Ebola virus killed 5,000 wild gorillas [12]. The severe population decline has contributed to the 2007 shift of the conservation status of western gorillas from “endangered” to “critically endangered” by the International Union for Conservation of Nature (IUCN) [14]. Furthermore, the recent outbreak of human EVD in North Kivu (Democratic Republic Congo) [3] was in close proximity to the remnant populations of eastern gorilla species, hence a human to ape transmission of the virus could potentially mark the end of existence for this critically endangered species.

Threats such as infectious diseases are relevant for conservation efforts, and efficient strategies are needed to reduce the effects of EVD on wild great ape populations [15]. Understanding any genetic impact that EBOV outbreaks might have on wild populations is vital, as EVD contributes to the fragmentation of gorilla populations due to a heterogeneous spatial influence of the outbreak [16]. Social dynamics in gorillas are rapidly affected by Ebola through a decrease in social cohesion, although recovery after the outbreak has been observed [17, 18]. One study reported that solitary individuals were less affected than individuals living in groups, marking the relevance of social dynamics for transmission [13]. A previous study using 17 microsatellites found no loss of genetic diversity after one EBOV outbreak in Lossi sanctuary and Odzala-Kokoua National Park, which could be explained by post-epidemic immigration, sufficiently large remnant effective population size or a short period of time after the decline [18]. The present study represents a continuation of this aforementioned research since many aspects of EBOV infection in wild gorilla populations are not yet explored, such as genetic variants in survivors that might contribute to resistance or higher chance of survival to EVD. In humans, such an approach was used during the 2014 EBOV outbreak, although no evidence of adaptation was found in the survivors [19].

Furthermore, microbial organisms inhabiting the gut also play a potentially crucial role in training and maintaining the immune system [20–22]. Where some commensal microbes are associated with priming the immune response or activating antiviral responses, others facilitate the development of the infection or suppress the immune response [23]. For instance, a recent study reported a link between the gastrointestinal microbiome of healthy humans and a predisposition to severe COVID-19 [24], with an abundance of Klebsiella, Streptococcus, and Ruminococcus being correlated with elevated levels of proinflammatory cytokine. The association between infectious diseases and the gut microbiome of nonhuman primates is less well understood. While the gut microbiome of SIVgor-infected (gorilla Simian Immunodeficiency virus) wild gorillas seems to be more robust to dysbiosis than those of chimpanzees and humans [25], it is unclear how shifts in the gorilla gut microbiome can impact the severity of viral infections, and in particular the immune response to Ebola virus infection.

Long-term monitoring of a western lowland gorilla (*Gorilla gorilla gorilla*) population in Odzala-Kokoua National Park (Republic of Congo) has been ongoing since 2001 until 2017 encompassing the Ebola outbreak of 2004 [16–18, 26] which resulted in a mortality rate of 95% [13]. Population monitoring involved the recording of individual histories of hundreds of identified individuals, the determination of sex, age and social status and the collection of fecal samples in different time periods. This close monitoring through time was fundamental to determine which individuals were either not affected or survived the Ebola outbreak and which went missing with their cause of death assumed to be EVD.

Here, we have obtained genetic data from gorilla fecal samples pre- and post-outbreak to explore potential associations to survivorship. Particularly, we compare the genetic variation at the single nucleotide level in 123 autosomal target-captured genes with putative roles in virus immune response and the gut microbiome composition in a reduced panel of survivors and non-survivors of this Zaire EBOV outbreak. With that we show that targeted capture on non-invasive fecal samples and next-generation sequencing can be used to study the impact of this severe disease in a natural population.

## Results

A total of 31 non-invasive fecal samples from identified western lowland gorillas were collected between 2001 and 2014 in Odzala-Kokoua National Park, Congo [18] (Fig. 1A). Sixteen of these were from individuals declared missing after the Zaire Ebola virus outbreak in 2004, and suspected to have died because of the infection, here termed ‘non-survivors’. These samples were collected between 2001 and 2004. The remaining 15 samples were from gorillas identified before the epidemic and still observed after the epidemic, and will be identified henceforth as ‘survivors’ [17]. These 15 samples were collected between 2005 and 2014 (Supplementary Table S1). We used target capture enrichment to sequence the genomic regions of 123 genes, which had previous evidence of putative roles in immune response to EBOV or other viruses (Supplementary Table S2). In addition, 15 neutral regions previously studied in other human and non-human primate studies were also targeted [27, 28] (Supplementary Table S2). Target design and all analyses were performed using the human reference genome due to the higher quality of annotations compared to the gorilla reference genome. We sequenced an average of 73 million paired reads per individual, 12% of which were unique (Supplementary Table S3). On average, 0.38% of the data mapped to the target space, representing an on-target effective coverage of 53.89-fold (range: 2.52-fold to 230.70-fold; Fig. 1B, Supplementary Table S3), with 72% of the target space covered by at least 4 reads per individual. Samples G282, G1392 and G638 performed poorly, with <50% of the target space covered at a minimum depth of 4 reads (Supplementary Fig. S1). Overall performance can be assessed by calculating how well the capture resequencing experiment went relative to expectations had we performed random shotgun sequencing. In that regard, we observe an average enrichment of 125-fold (88 - 346-fold) (Supplementary Table S3). Individuals with extremely low proportions of target space covered by at least 4 reads (<30%) (G638 and G282) and high heterozygosity and high levels of human contamination were removed from further analysis (individuals G374 and G1392, Fig. 1C and Supplementary Fig. S2, Table S3).

**Fig. 1.**
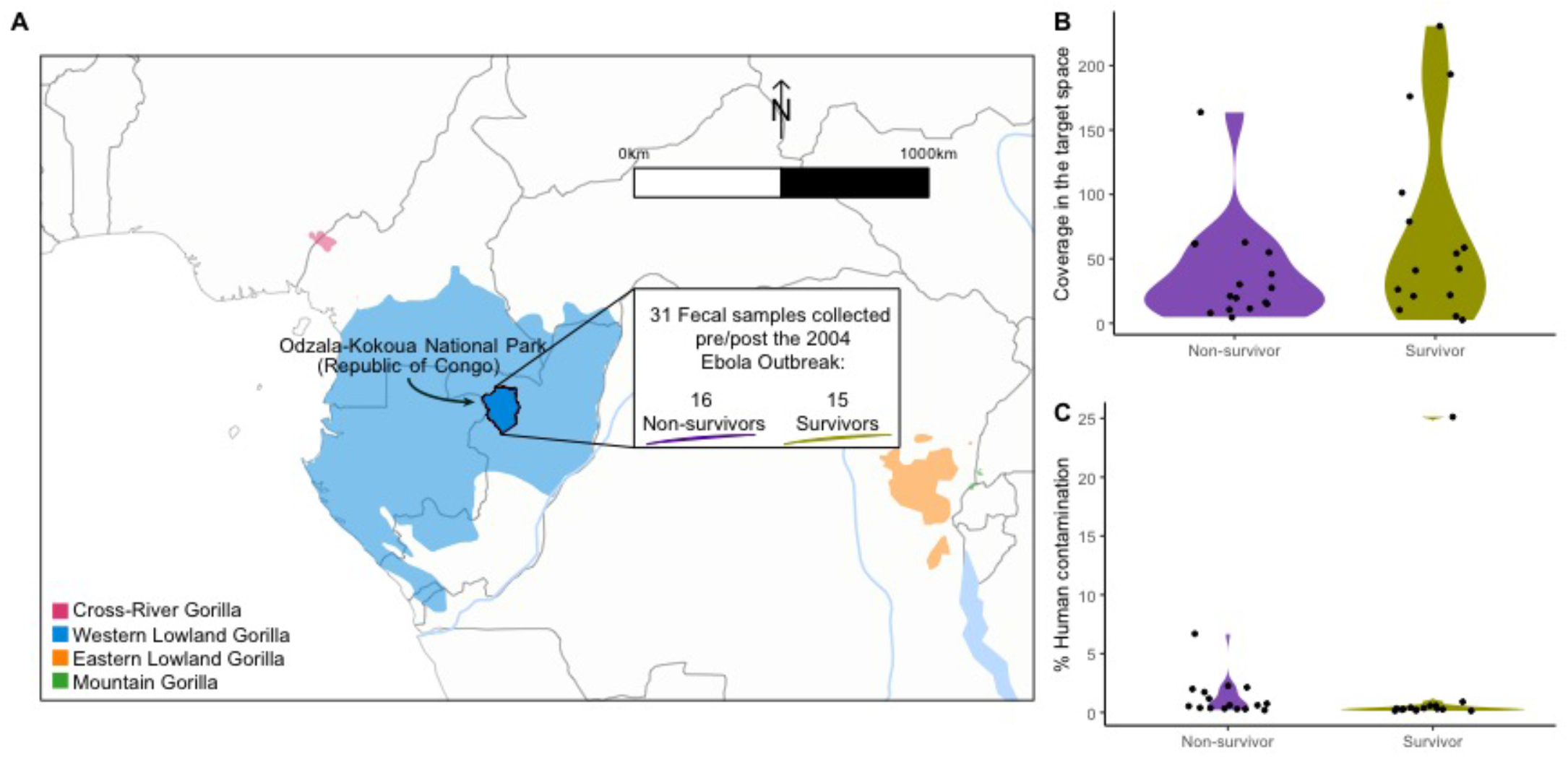
Sample description. **A**) Geographical map of the extant range of gorillas and the Odzala-Kokoua National Park (Republic of Congo) where fecal samples were collected between 2001 and 2014, overlapping the Ebola outbreak in 2004. **B**) Average coverage reached in the target space per sample in both studied groups. **C**) Percentage of human contamination in each fecal sample.

With the final dataset of 13 survivors and 14 non-survivors, we validated that the genotype information obtained was in concordance with previously published gorilla whole-genomes, as determined by a principal component analysis (PCA) (Supplementary Fig. S3). We estimated kinship for each pair of individuals, identifying a single case of close relatedness (1^st^ degree) among them (non-survivor G374 and survivor G739; kinship coefficient = 0.4) (Supplementary Fig. S4). In addition, we observed no stratification correlating with survivor/non-survivor classification. Individuals appear to be dispersed randomly across a dendrogram derived from shared genotype likelihood dosage states (Fig. 2A), and a univariate linear regression of survivorship on the top 5 PCs identified no significant structure associated to survivorship (P-value > 0.1; Supplementary Fig. S5). Hence, we determined no genome-wide group structure differences between survivors and non-survivors. The overall level of genetic diversity within the target space of the studied gorillas was, on average, lower than that of western lowland gorillas obtained from whole-genome sequencing (Supplementary Fig. S6) [29, 30], an expected outcome following the target capture procedure. Moreover, there are no statistically significant differences in heterozygosity between survivors and non-survivors (Student’s t-test, p-value=0.34; Fig. 2B).

**Fig. 2.**
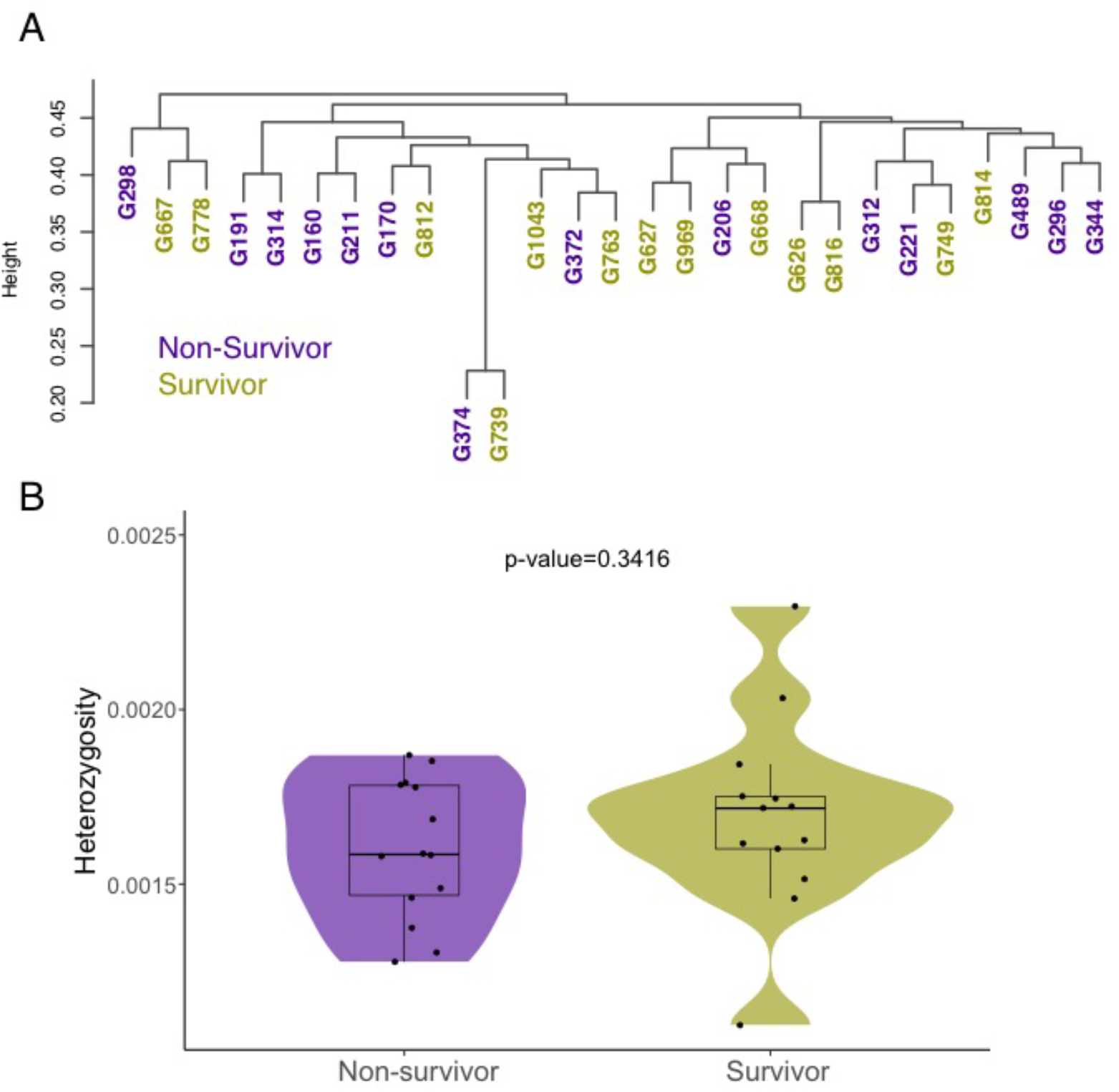
Genetic distance dendrogram and heterozygosity among non-survivor and survivor gorilla groups. **A)** Clustering dendrogram of pairwise genetic distance derived from genotype likelihoods (N=5,477). **B)** Mean heterozygosity (bp^-1^) in non-survivors and survivors; not significantly different (Student t-test, p-value=0.34).

In order to determine genetic differences between the groups, we calculated three summary statistics on a dataset of 6,852 high-quality variants: (1) the difference in allele frequency (ΔFrequency), (2) the fixation index (*F*_*ST*_), and (3) the significance level (α) of each variant for its association with the binary trait survivor/non-survivor (Fig. 3). We found 118 SNPs within the target space that surpassed our α threshold in the association test, and we also reported their ΔFrequency and *F*_*ST*_ values. However, after controlling for type I error none of these remained significant. Out of these, seven genes have multiple nominally significant SNPs (*CD1B, IGKV4-1, HLA-A, ACTB, LYN, CD68* and *MX1*), while 10 neutral regions (∼10kb each) have at least 1 nominally significant variant (Supplementary Table S4). For comparative validation, we repeated the association analysis using ANGSD [31], a software explicitly built to work with low coverage data that relies on genotype likelihoods. With this method, we were able to recover the majority of the genes found above (30 out of 36). However, ANGSD returned more hits and thus more genes (Supplementary Figs. S7, S8 and Table S5), rendering the above approach more conservative.

**Fig. 3.**
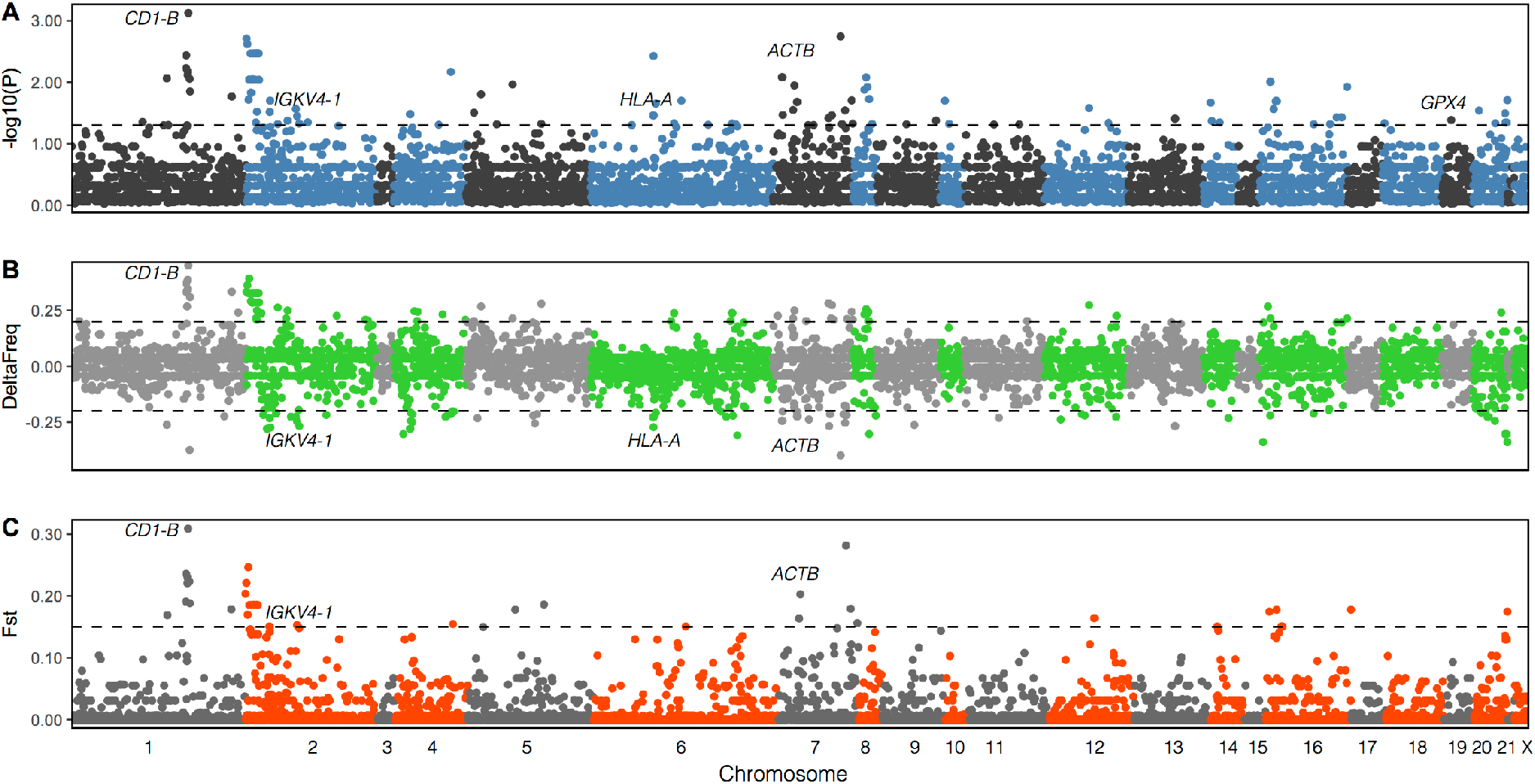
Association analysis to detect SNPs and candidate genes related to survivorship to EBOV outbreak. **A)** Significance level (threshold set at α=0.05, -log10(P-value)=1.30). **B)** Difference in allele frequency (threshold set at ±0.2). **C)** Fixation index (threshold set at *F*_*ST*_=0.15). Dashed lines delineate the thresholds used.

Next, we explored the potential functional impact of the variants in the nominally significant candidate loci that differentiate survivors from non-survivors. While we found no significant associations with gene ontology categories, the analysis of predictions of functional consequences pinpointed six missense mutations which differed in frequency between the two groups (Supplementary Table S6), one each in the *ATM, IGKV4-1* and *RNF167* genes (all lower in survivors) and three in the *ACTB* (Actin Beta) gene (one unique to survivors, two lower in survivors, Supplementary Table S4). All three missense variants in *ACTB* are predicted as deleterious by both the PolyPhen [32] and SIFT [33] algorithms. Furthermore, the derived variant in survivors in the immunoglobuline-encoding *IGKV4-1* might be deleterious (C-score > 20 [34]), hence potentially functionally relevant. Since we used the human genome for target design, mapping and variant calling, we caution that differences in exon usage or pseudogenization on the gorilla lineage might confound these inferences of protein-coding changes. In order to confirm the expression of these genes, and specifically the exons of interest, we mapped transcriptome data from six tissues in gorillas [35] to the same reference genome, and quantified expression levels. We confirmed the expression of these genes and found high transcript abundance (log2-value of counts >=9) for *ACTB, RNF167* and *ATM*, while *IGKV4-1* was only detected at low levels in these tissues (log2-value of counts = 5.84). We found ∼10,000 RNA sequencing reads overlapping the three loci of interest in *ACTB*, 437 in *RNF167*, and 96 in *IGKV4-1*, supporting the expression of these specific loci in gorilla tissues, while for *ATM* only 10 reads overlapped (Supplementary Table S6).

Among the 118 significant SNPs, we found no direct overlap with loci associated to 2,385 traits in genome-wide association studies (GWAS) for humans [36]. However, we found 49 associated loci within close proximity (5,000 bp) to GWAS loci (Supplementary Table S7), among which the locus 19:1104078 near GPX4 stands out for its association with five blood cell traits (counts and percentages of leukocyte cell types). Furthermore, 6:29916885 in the *HLA* is associated to hemoglobin levels. Five loci in *HLA-DRA* are associated with different autoimmune diseases like systemic lupus erythematosus or multiple sclerosis (Supplementary Table S7). Furthermore, nine loci in this region (*HLA-DRB5* and *HLA-A*) are associated with schizophrenia or autism spectrum disorder (Supplementary Table S7).

It has been reported that intestinal microbiota play a critical role in immune response to infectious diseases [20–22]. Thus, the microbiome might be relevant for the survival of wild gorilla populations experiencing an Ebola outbreak. Taking advantage of the nature of the samples, we first analyzed the microbiota present in each fecal extraction using a 16S rRNA library (Methods). We obtained a total of 96,928,400 reads with an average sequencing depth of 1,101,459 (SD ± 418,989) reads per gorilla fecal extraction (Supplementary Table S8), and determined the abundance of taxa (Supplementary Figs. S9 and S10). Firmicutes (53.79%), Bacteroidetes (12.02%) and Chloroflexi (11.11%) were the predominant phyla (Supplementary Fig. S9 and Table S9), that include the following most abundant orders: Clostridiales (39.37%), Bacillales (11.24%), Bacteroidales (10.87%) and Anaerolineales (11.11%) (Supplementary Fig. S10 and Table S10), concordant with previous findings [37, 38]. We found no taxa significantly differing in relative abundance between survivor and non-survivor gorillas (Bonferroni-corrected p-values > 0.05; Supplementary Fig. S11 and Table S11), and sample groups were not separated in a clustering analysis (Fig. S12).

Since these results on the gut microbiome diversity did not support differences between both gorilla groups, we decided to perform deep sequencing on the fecal libraries. We generated a total of 801,132,281 sequences from DNA libraries (4,025,054-25,593,317 reads per sample; Supplementary Table S8) and used MALT (MEGAN Alignment Tool) [39] for an alternative characterization of the microbial profile of the samples (Supplementary Fig. S13). We find that the majority of the identified taxa are associated with the gut microbiome (Supplementary Table S12 and S13). By far the most abundant taxon is the gut bacterium *Escherichia coli*, which could be detected in all samples, and makes up more than 50% of all assigned sequences in nine of the samples. Also in high abundance are species of the Bacteroidales order, such as *Bacteroides cellulosilyticus* and *Prevotella spp*., as well as members of the Clostridiales, Lactobacillales, and Bacillales orders, corroborating previous reports on the composition of western lowland gorilla gut microbiomes [40, 41]. Furthermore, we detected pathogenic taxa in high abundance in some of the samples, such as *Clostridium botulinum, Acinetobacter baumannii*, and *Klebsiella pneumoniae*, which have been previously found in the gorilla gut [42]. However, the microbial profiles of survivors and non-survivors do not differ significantly from each other in this analysis either (Bonferroni-corrected p-values > 0.05, two-sided t-test, Supplementary Table S11), and the two groups do not form separate clusters in a Principal Coordinate analysis or a Neighbor Joining Tree (Supplementary Figs. S14 and S15).

## Discussion

We investigated non-invasive fecal samples from a long-term monitored population of western lowland gorillas in the Republic of Congo, including individuals that most likely succumbed to the *Zaire* Ebola virus outbreak in 2004, as well as surviving individuals [17, 18]. We used targeted capture of 123 autosomal genes with putative roles in immune response to EBOV or other viruses (Supplementary Table S2) from fecal samples. This yielded an enrichment of more than 100-fold across samples, and a medium to high coverage of the target space across most individuals (Supplementary Fig. S1 and Table S3). Although a large proportion of reads were duplicates, the overall performance was high and these results demonstrate the great potential of capture experiments for obtaining genotypes from fecal samples of wild great ape populations [43, 44], for which high-coverage sequencing would be prohibitively expensive. We determined that the studied individuals were not closely related, hence most likely representing a random sampling of the wild gorilla population before and after the outbreak. We also investigated the microbial community composition of survivors and non-survivors, finding no significant differences in taxa abundance, neither using 16S rRNA or deep sequencing data (Supplementary Figs. S11, S13 and Table S11). Hence, we find no evidence that the gut microbiome of individuals has an influence on the survival rate of wild gorillas exposed to Ebola. However, these observations are limited by (1) the sample size, and (2) the broad range of collection dates (Supplementary Table 1). The latter is particularly true (2a) relative to the timing of any exposure, but also in respect to the (2b) dynamic nature of the gut microbiome [38].

Given the limited sample size, we developed an approach using differences in allele frequency, the fixation index and the effect size to determine variants most strongly associated to survivability in the studied population, generally replicable using an association analysis with ANGSD. While 44 of the 118 nominally significant SNPs (Supplementary Table S4) do fall within 10 of the 15 neutral regions included in the study, some SNPs might be functionally relevant for surviving the EBOV outbreak. The non-synonymous variants in *ACTB, RNF167* and *IGKV4-1* genes are obvious candidate loci, and particularly the three deleterious missense mutations in Actin Beta appear to be strong candidates for a higher survival rate. The actin cytoskeleton is important for virus assembly [45], and a disturbed assembly process could have influenced the viral load in individuals with changes in this protein. As expected for a gene encoding a structural protein, *ACTB* is highly expressed in gorilla tissues. Furthermore, the variant in *IGKV4-1* might improve the immune response to viral infection through antigen recognition [46]. The missense mutation in ATM, which belongs to the PI3-kinase family, could interfere with the cellular entry specifically of the Ebola virus [47], although we could not confirm expression of this locus *in vivo* in the available tissues.

We find other potentially relevant non-coding variants, 49 of which are in close proximity to SNPs associated to GWAS traits in humans, suggesting possible regulatory functions (Supplementary Table S7). Among those, the association of *GPX4* with leukocyte cell type count might reflect differences in leukocyte composition after viral infection. Differences in hematocrit or hemoglobin levels might have contributed to the survival of wild gorillas considering that hemorrhage and internal bleeding are symptoms of Ebola infection. Since eight loci are associated to the *HLA-DRB* gene, a direct involvement in the adaptive immune system might cause the signature observed at this locus, particularly given that human survivors of EVD show a lower frequency of *HLA-DR*-positive T cells [48].

## Conclusion

By using fecal samples and targeted capture enrichment, non-invasive assessment of numerous individuals from wild populations is possible. Here, we demonstrate that this approach can be used to analyze temporal genetic changes in wild great ape populations in response to environmental factors. Additionally, we present candidate loci that may have facilitated the survival of gorilla individuals or groups after an outbreak of the *Zaire* Ebola virus. Understanding putative adaptive responses to this pathogen in wild populations can help to advance our knowledge on the natural dynamics of this severe disease. Such a strategy might be useful in a broader context, since these and other primates are susceptible to other infectious diseases such as Covid-19 [49].

## Methods

### Samples, DNA extraction, library preparation

Non-invasive fecal samples from western lowland gorillas were collected between 2001 and 2014 in Odzala-Kokoua National Park, Republic of Congo [18]. Among them, we selected 31 samples from previously identified individuals. Sixteen of those individuals were declared missing after the epidemic in 2004 [13]. They died during the time span of the epidemic and were identified here as non-survivors. Fifteen individuals were observed before and after the epidemic, and were described here as survivors (Fig. 1A and Supplementary Table S1) [17]. Samples were collected by field investigators wearing masks and gloves, and were dried with silica beads and then stored at room temperature until arrival in a laboratory where they were stored at 4°C until extraction.

DNA was extracted from 10mg of dried sample using the 2CTAB/PCI protocol [50] using negative controls that were checked for contamination before subsequent experiment. Three different extractions were carried out, except for samples G778, G374, G344, G498 and G372, where only two extractions were performed (Supplementary Table S1). A DNA library [51–53] and a 16S rRNA library [54–56] were prepared for each extract. Isolated DNA samples were quantified with Qubit with a mean estimated concentration of 13.3 ng/µl (range: 0.90-74.7). Whenever possible, a total of 250 ng of DNA was used to construct DNA libraries, but never more than a total volume of 33 µl was taken from any single sample. DNA was sheared with a Covaris S2 instrument and 88 fecal DNA (fDNA) libraries were prepared following a custom dual-indexing protocol with 25 cycles of amplification [51, 52]. Subsequent to DNA library preparation, 88 16S rRNA libraries were prepared using 1 µl of total DNA. The V3 and V4 regions of 16S rRNA were target amplified using modified 341F and 806Rb primers [54, 55, 57], incorporated into the dual-indexing protocol [52]. The forward primer (IS1_P5_16S_341f: ACACTCTTTCCCTACACGACGCTCTTCCGATCTNNNNCCTACGGGNGGCWGCAG), and reverse primer (IS2_P7_16S_806rB: GTGACTGGAGTTCAGACGTGTGCTCTTCCGATCTGGACTACNVGGGTWTCTAAT) include the complementary sequences necessary for the final indexing step [52]. Protocols are provided in [53].

### Target Design, Capture and Sequencing

RNA baits covering the target space were designed and synthesized by Agilent with a minimum of 3x bait coverage. The target space included specific autosomal genes (123 genes) and 15 neutral regions (∼10kb each) (Supplementary Table S2). For target enrichment, the fDNA libraries were pooled into one equimolar batch and subjected to two consecutive rounds of DNA capture with the RNA baits in 8 hybridizations. Captured fDNA libraries were sequenced on the Illumina system in four HiSeq 2500 2×125 lanes and one HiSeq 2500 rapid run at 2×250 bp. The 16S libraries were sequenced on one HiSeq 2500 rapid run 2×250 bp lane. In addition, we generated paired-end sequences from the fDNA libraries on four HiSeq 4000 lanes (2×150bp) to study the whole microbiome composition (Table S8).

### Mapping and Variant Discovery

Prior to mapping, paired-end reads belonging to the same library but sequenced in different lanes were merged into a single FASTQ file. PCR duplicates were directly removed from FASTQ files using FASTuniq (v1.1) [58]. Overlapping reads were merged (minimum overlap of 10 bp, minimum length of final read to 50 bp) using PEAR (v0.9.6) [59]. Reads were mapped using BWA mem (v0.7.12) [60] to the human reference genome Hg19 (GRCh37 from the UCSC database). Assembled reads were mapped considering single-end specifications and unassembled reads considering paired-end specifications. Any remaining PCR duplicates were removed using PicardTools MarkDuplicates (v1.95) (http://broadinstitute.github.io/picard/). Non-primary alignments and reads with quality below 30 were filtered from the dataset with samtools (v1.5) [61]. Finally, single-end and paired-end reads were merged into a single BAM file using PicardTools MergeSamFiles (http://broadinstitute.github.io/picard/). The percentage of aligned reads for each DNA extraction and sample was calculated by dividing the number of uniquely and high-quality mapped reads (without duplicates) by the total number of sequenced reads. The percentage of on-target aligned reads was calculated for each sample by dividing the number of on-target filtered reads by the number of sequenced reads. The average target effective coverage was calculated dividing the number of aligned bases by the total length of the targeted genomic space. Finally, the enrichment factor (ER) of the capture performance was calculated using the ratio between the on-target reads by the total mapped reads over the targeted size by genomic size (ER = (On-Target Reads/Mapped Reads)/(Target Size/Genome Size)). The coverage for each target region was retrieved using SAMtools bedcov [61].

For variant calling, all BAM files belonging to the same sample were merged into a single BAM file using PicardTools MergeSamFiles (v1.95) (http://broadinstitute.github.io/picard/). Variant discovery was performed using GATK ‘Unified Genotyper’ [62] for each sample independently with the following parameters -out_mode EMIT_ALL_SITES -stand_call_conf 5.0 -stand_emit_conf 5.0 -A BaseCounts -A GCContent -A RMSMappingQuality -A BaseQualityRankSumTest. Afterwards, we merged each sample gvcf to a single one using GATK ‘CombineVariants’ [62] with the following parameters - genotypeMergeOptions UNIQUIFY –excludeNonVariant. We also included in the gvcf the genotype information of available whole genome data of six *Gorilla beringei beringei*, eight *Gorilla beringei graueri*, one *Gorilla gorilla dielhi*, and twenty-three *Gorilla gorilla gorilla* samples [29, 30]. The VCF was filtered with VCFtools [63] to keep only biallelic positions with DP >3 and quality > 30 and without indels.

Genotype likelihoods were directly obtained from BAM files with ANGSD [31] including four *Gorilla beringei beringei*, four *Gorilla beringei graueri*, one *Gorilla gorilla dielhi*, and four *Gorilla gorilla gorilla*, with the following parameters and only in the target space: -uniqueOnly 1 -remove_bads 1 -only_proper_pairs 1 -trim 0 -C 50 -baq 1 -minInd 21 -skipTriallelic 1 -GL 2 -minMapQ 30 -doGlf 2 -doMajorMinor 1 -doMaf 2 -minMaf 0.05 -SNP_pval 1e-6.

### Quality control

We evaluated the amount of human contamination in each fecal library using the HuConTest script [64], as described previously [65]. The majority of samples have less than 2% of human contamination, but samples G348 and G1392 have estimates of human contamination of 6.7% and 25.1%, respectively (Supplementary Table S3). These two samples also show extreme values of heterozygosity (deviating >1s.d. from mean heterozygosity; Figure S2). For the identification of individuals and markers with elevated missing data rates we used the proportion of the target space covered by at least 4 reads. Individuals with less than 30% of covered target space (4 reads) were not used for further analysis (G638 and G282).

A principal component analysis (PCA) was performed to validate that the genotype information obtained for the case study gorillas was in concordance with previously published data. We used PCAngsd [66] with the genotype likelihoods obtained with ANGSD (N=6,484), including 13 previously published whole-genomes representative of each know gorilla subspecies [29, 30]. We also obtained a PCA using the GATK genotype calls after keeping only variants with minor allele frequency of 0.02 with plink --pca option (N=6051) [67].

### Genetic distance, relatedness and heterozygosity

We used the genotype likelihood information for the studied individuals to obtain the genetic distance by running ngsDist in ANGSD [68] with the following parameters: --n_sites 5477 --probs TRUE --pairwise_del. Then, we constructed an Euclidean distance matrix based on the genotypes and performed a hierarchical clustering using the R package *ape* [69]. We also run PCAngsd [66] considering only the study gorillas to discard any possible intra-group structure.

The theta coefficients of kinship (probability of a pair of randomly sampled homologous alleles are identical by descent) were calculated using the NgsrelateV2 [70, 71] on the genotype likelihood obtained with ANGSD [31]. Note that all possible genotype likelihoods, even outside the target space (N=226,094), were used since the coverage of the kinship markers was insufficient.

To assess global levels of heterozygosity, the unfolded SFS was calculated for each sample separately, including thirteen gorilla whole-genomes representative of all gorilla subspecies [29, 30], using ANGSD [31] and realSFS [72] only in the target space with the following quality filter parameters: -uniqueOnly 1 -remove_bads 1 -only_proper_pairs 1 -trim 0 -C 50 -baq 1 -minMapQ 20 -minQ 20 -setMaxDepth 200 -doCounts 1 -GL 1 -doSaf 1. We used the human genome (Hg19) to determine the ancestral state.

### Association analysis

The genotype calls obtained with GATK were further filtered with Plink [67] to exclude variants considering their missing rate (--geno 0.05), minor allele frequency (--maf 0.01) and Hardy-Weinberg equilibrium (--hwe 0.00001). The final dataset consists of 27 samples (13 survivors and 14 non-survivors) and 6,852 high-quality variants. Nominal significance was set at an alpha of 0.05 and a Bonferroni corrected alpha threshold of 7.3×10-6 (0.05/6,852) was defined to account for familywise error. Associations were tested for by a chi-square allelic test with one degree of freedom and p-values were estimated by permutation in plink (plink –assoc –mperm 10000). [73]. The p-values were plotted in a Manhattan plot in R (v3.4.1).

The allele frequency for each group (survivors and non-survivors) was obtained using the -freq2 option in VCFtools [63]. Then, we calculated the allele frequency difference per SNP by subtracting the allele frequency in non-survivors from the allele frequency in survivors (ΔFrequency). We chose a threshold of ± 0.2, and plotted the allele frequencies using R. The fixation index (*F*_*ST*_) between both groups was calculated using VCFtools –weir-fst-pop option (Weir and Cockerham) [63] with a threshold at 0.15, and results were plotted in R. We retrieved markers with α ≤ 0.05 in the association test and a p-value < 0.05 in the permutation test. The ANGSD software [31] was used to perform a replication of the association analysis using the following parameters -minQ 20 -minMapQ 30 -doAsso 1 -GL 1 -out assocGQ_filter - doMajorMinor 1 -doMaf 1 -SNP_pval 1e-6 -minInd 22 -minMaf 0.02. The output of the association analysis are LRT values (Likelihood Ratio Test), which are chi square distributed with one degree of freedom. Since we set a threshold of significance at 95% confidence, the minimum score to be significant is LRT = 3.84. In both association analyses, we linked the nominally significant SNPs with their genes (Supplementary Table S4 and S5). Genes with multiple nominally significant SNPs were considered to be potentially more relevant. Subsequently, we compared the overlap of discovered genes between the datasets (Unified Genotyper and ANGSD) in a Venn diagram (Supplementary Fig. S8) using the R package VennDiagram [74].

### Prediction of functional consequences of the significant markers

We used VEP (v91) [75] for the functional annotation of the associated SNPs. We retrieved the predicted consequence of each significant marker found in the potentially related genes to Ebola immune response, as well as PolyPhen-2 [32], Sift [76] and C-scores [77]. Associated loci were intersected with hits in the GWAS catalogue [78] within 5,000 bp. We also performed an overrepresentation test using Panther [79] to test whether any of the potentially related genes are overrepresented in biological or functional categories compared to the rest of targeted genes with no apparent association with Ebola. In addition, we mapped previously published RNA sequencing data from six tissues (brain, cerebellum, heart, kidney, liver, testis) in two gorilla individuals [35] to the annotated genes in the human reference genome (using the Ensembl Release 75 gene models) using Tophat2 [80], and estimated the gene expression with htseq-count [81]. Gene expression is reported log2-normalized, and we counted the number of reads overlapping the candidate missense mutations to confirm their transcriptional activity in gorillas. Values presented are the cumulative sums of RNA sequencing reads across individuals and tissues.

### Microbiome sequencing

16S RNA sequencing reads were processed using QIIME (v1) (Quantitative Insights Into Microbial Ecology) [82] to analyze the 16S rRNA. First, paired-end raw reads were merged using fastq-join from ea-utils package [83]. Then, with usearch software [84], merged FASTQ reads were filtered (-fastq_trunclen 253 and –fastq_maxee 0.5). Using QIIME environment, the metadata mapping file was constructed and validated (validate_mapping_file.py) and QIIME labels were added (add_qiime_labels.py). We applied open-reference OTUs picking (pick_open_reference_otus.py). Summary statistics were computed using *biom summarize-table*. The resulting dataset was rarefied to an even depth of 10,000 sequences per extract (6 extracts were excluded in diversity analysis: G191_5782, G344_5827, G314_5824, G489_5834, G489_5835, G344_5828). Finally, we ran diversity analysis with a sequence depth of 10,000 (core_diversity_analysis.py). Taxa abundance quantification and significance or relative taxa abundance (T-test and p-values adjusted for multiple testing with Bonferroni-correction) were computed in R.

Deep sequencing of the DNA library (pre-capture) was performed as stated above. To remove sequencing adapters and merge the read pairs, we used AdapterRemoval v2.2.4 with default settings [85]. We then aligned the merged sequences to the gorilla reference genome Kamilah_GGO_v0 using bwa mem [60] to remove host DNA. Subsequently we filtered out potential human contaminant DNA by aligning the unmapped sequences to the human reference genome hg19, resulting in 724,738,878 filtered sequences. MALT v0.4.1 (MEGAN Alignment Tool) [39] was used to characterize the microbial profile, using all archaeal, viral, and bacterial reference sequences downloaded from NCBI on 06.05.2019. These were indexed using malt-build to build a custom database. Malt-run was then used with minimum percent identity (--minPercentIdentity) set to 95, the minimum support (--minSupport) parameter set to 10, and the top percent value (--topPercent) set as 1, other parameters were set to default. The resulting rma6 files were visualized with MEGAN6 [86] and clustered in a Principal Coordinate analysis (PCoA) and Neighbor Joining Tree analysis according to microbial composition on the species level (Supplementary Figs. S14 and S15).

## Availability of data and materials

The dataset generated during the current study will be made publicly available upon acceptance (raw sequencing data to ENA, project ID: PRJEB43265). It will be available to reviewers from the corresponding author on request.

## Abbreviations

EBOV: Ebola virus
EVD: Ebola virus Disease
GWAS: Genome-Wide Association Study
IUCN: International Union for Conservation of Nature
PCA: Principal Component Analysis
SNP: Single Nucleotide Polymorphism

## Declarations

## Acknowledgments

We are grateful to S. Gatti, F. Levréro, C. Genton, R. Cristescu for their assistance with sample collection on site. We thank the teams of the ECOFAC program (EU) and African Parks Networks for logistic assistance and permission to work in Odzala-Kokoua National Park. The storage and extraction of fecal samples were performed in the molecular ecology platform (UMR 6553 Ecobio, CNRS/UR1) dedicated to non-invasive samples.

## Contributions

D.H., T.M.-B., P.L.G., N.M., D.V., A.N. and M.T.P.G. conceived and designed the study; D.H., D.V. and C.S.O. performed experiments; C.F., M.K., D.H., J.H.R., C.H.S.O and J.N. analyzed data; P.L.G., N.M. and D.V. collected genetic and demographic data; P.F., H.R.S., C.H., provided analytical support. C.F., M.K., D.H., and T.M.-B. wrote the manuscript with input and approval from the other authors.

## Corresponding authors

Correspondence to: Claudia Fontsere, Tomas Marques-Bonet or Martin Kuhlwilm

## Ethics declarations

### Ethics approval and consent to participate

Not applicable.

### Consent for publication

Not applicable.

## Additional information

**Additional File 1**. Excel file with tables S1-S13.

**Additional File 2**. PDF with Figures S1-S15.

