## Supplemental Figures for "The genetic impact of an Ebola outbreak on a wild gorilla population"

### Supplementary Figures S1-16

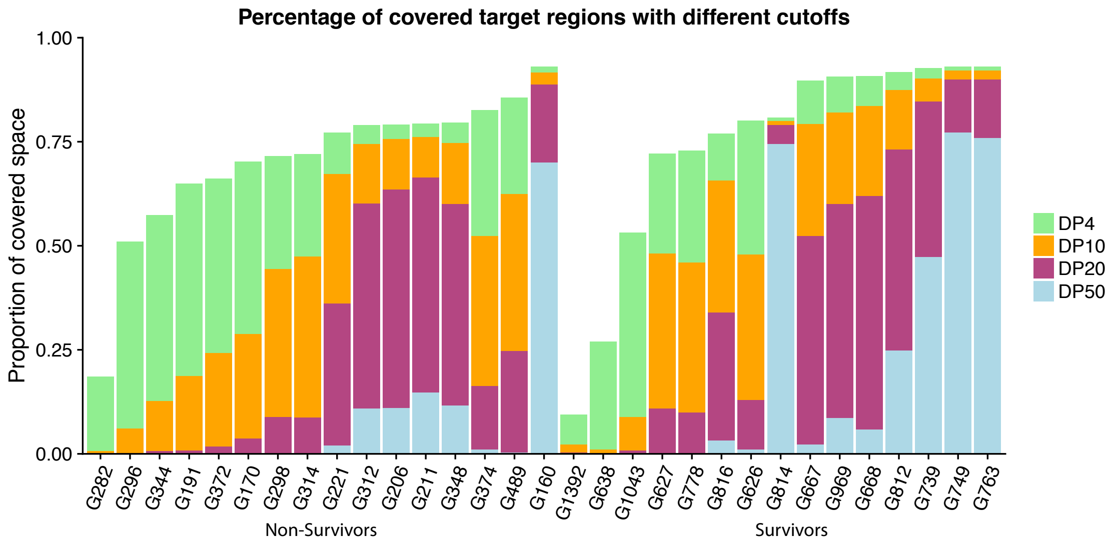


Fig. S1. Percentage of covered target regions considering different depth of coverage (4x, 10x, 20x and 50x).

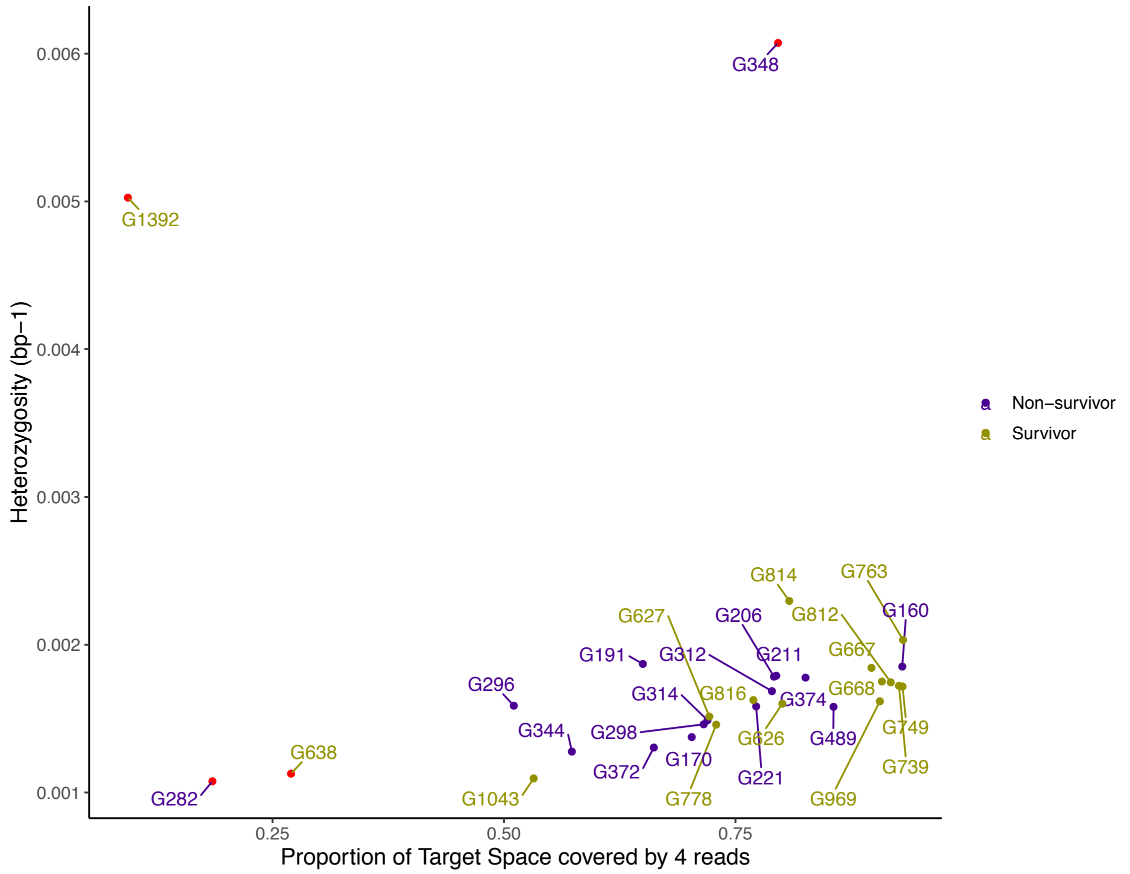


Fig. S2. Proportion of the target space covered by at least 4 reads with average heterozygosity of each sample. Two samples exhibit higher than expected heterozygosity (G348, G1392) and other two samples have very low coverage with a less than 30% of the target space covered (G638 and G282). These outliers are colored in red.

A

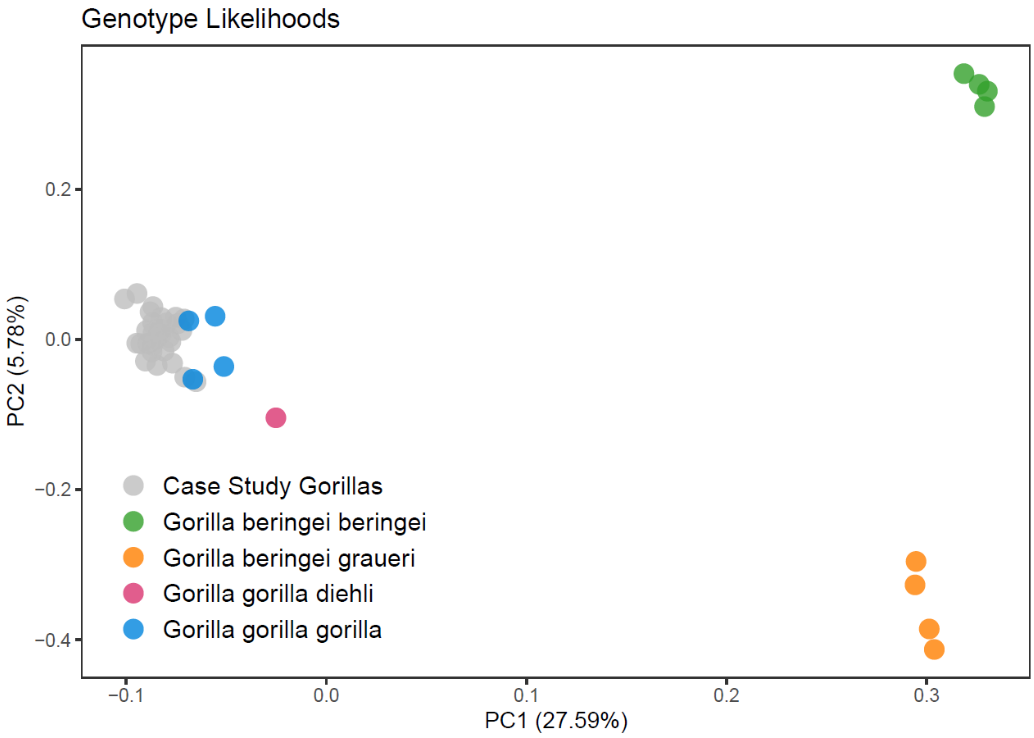


B

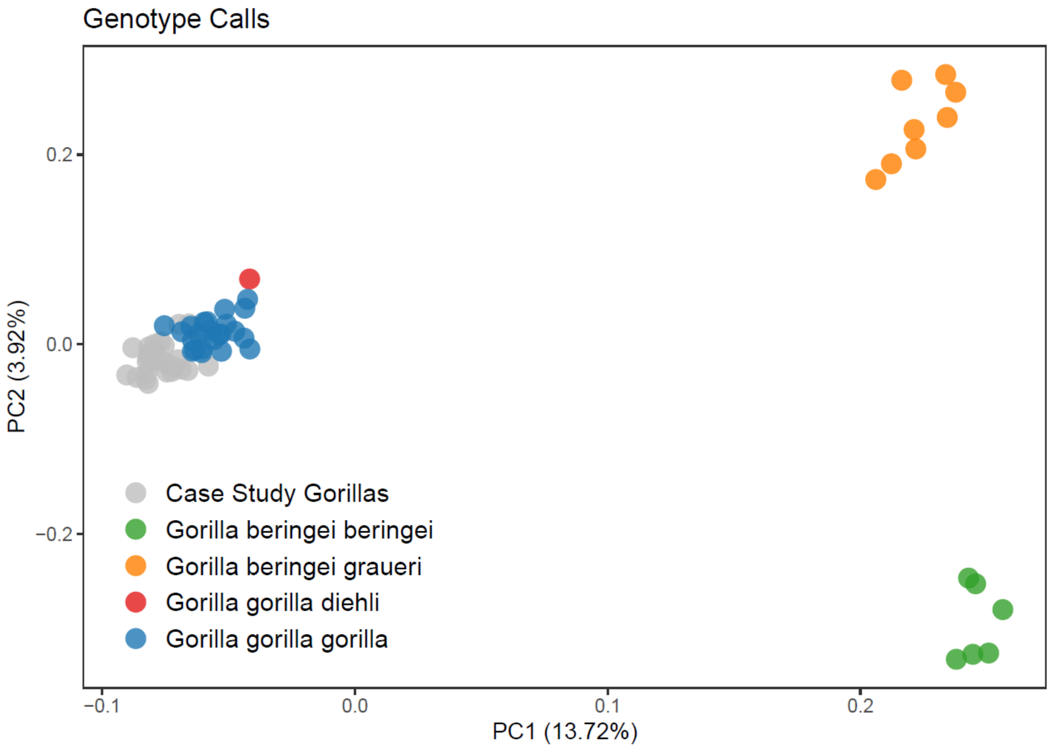


Fig. S3. Principal Component Analysis (PCA) of previously published gorilla genomic data (Prado-Martinez et al 2013, Xue et al 2015) together with the case study gorillas. A) Using genotype likelihoods obtained with ANGSD from 13 available bam files from published genomes (four per subspecies except Cross

River Gorillas, for which only one exists; 6,484 variants) and B) Using genotype calls obtained with GATK from 38 published genomes (6,051 variants), the largest dataset on gorilla variation, as described elsewhere (Kuhlwilm et al., 2021).

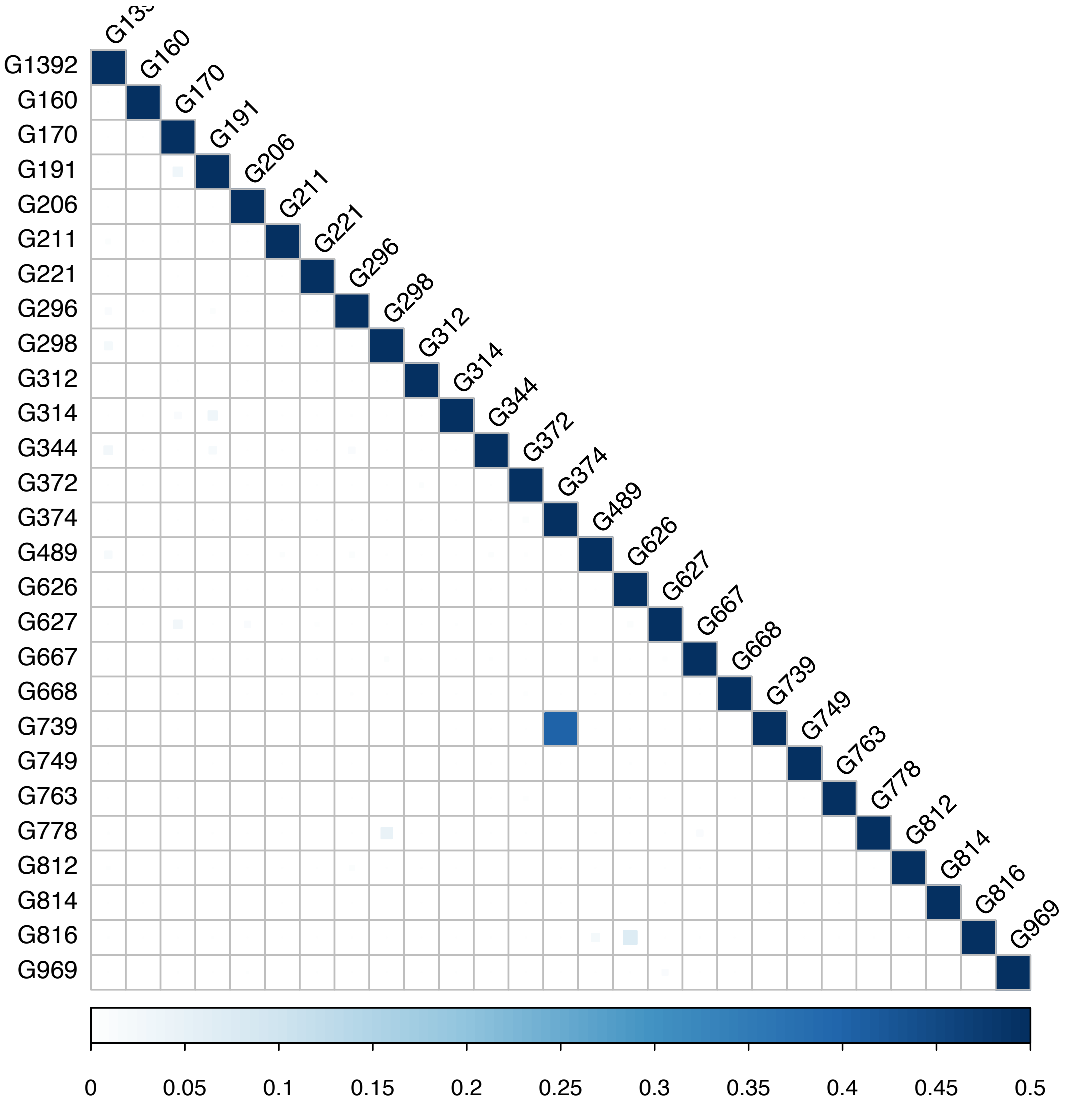


Fig. S4. Pairwise relatedness estimates for the case study gorillas derived from genotype likelihoods (N=226,094). Only one pair of samples (G739-G374) is found to be 1st order relatives (kinship coefficient 0.4).

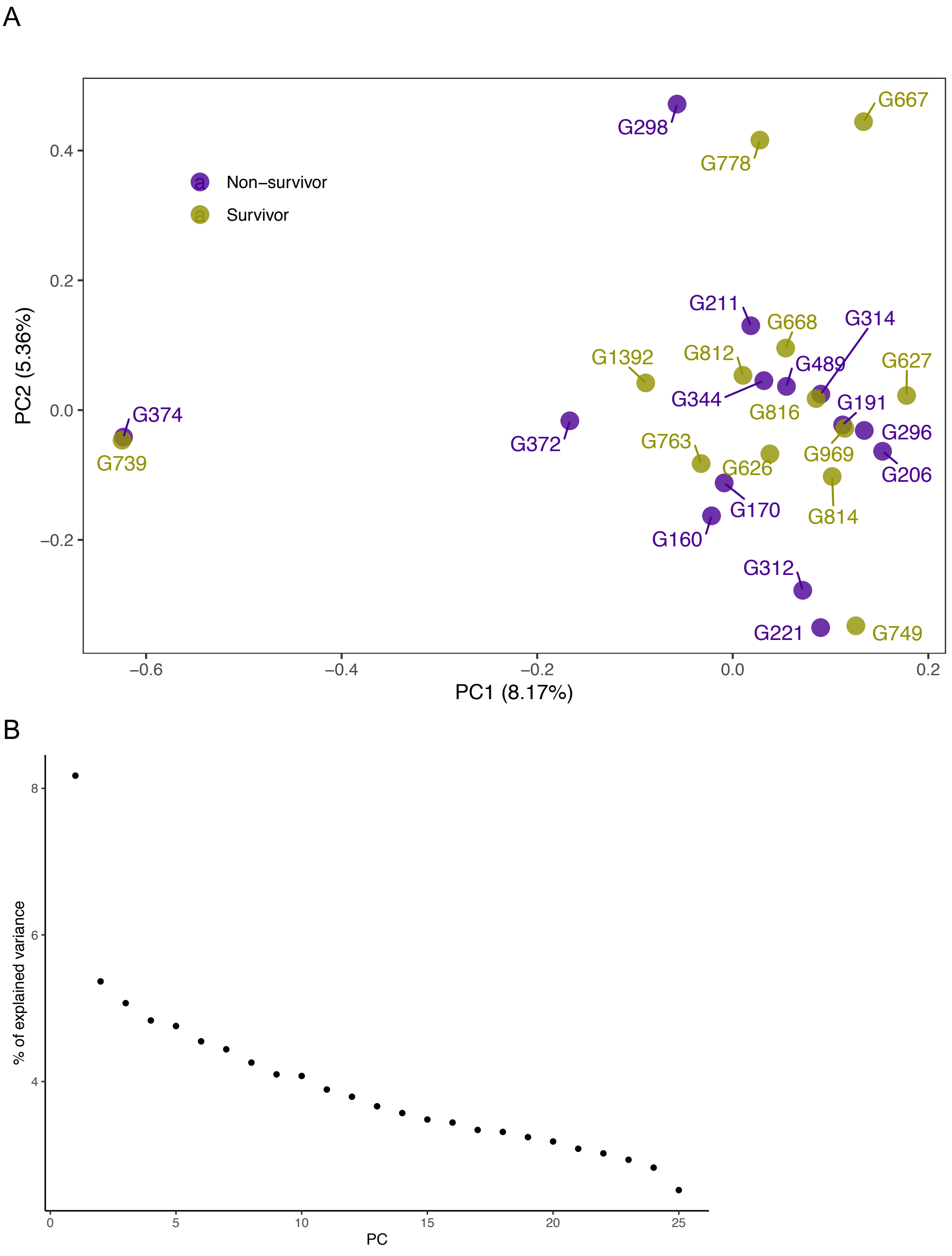


Fig. S5. Principal Component Analysis (PCA) of the case study gorillas derived from genotype likelihoods from ANGSD (5,833 variants). A) PC1 vs PC2 and B) Scree plot with the % of the variance explained for 25 PCs.

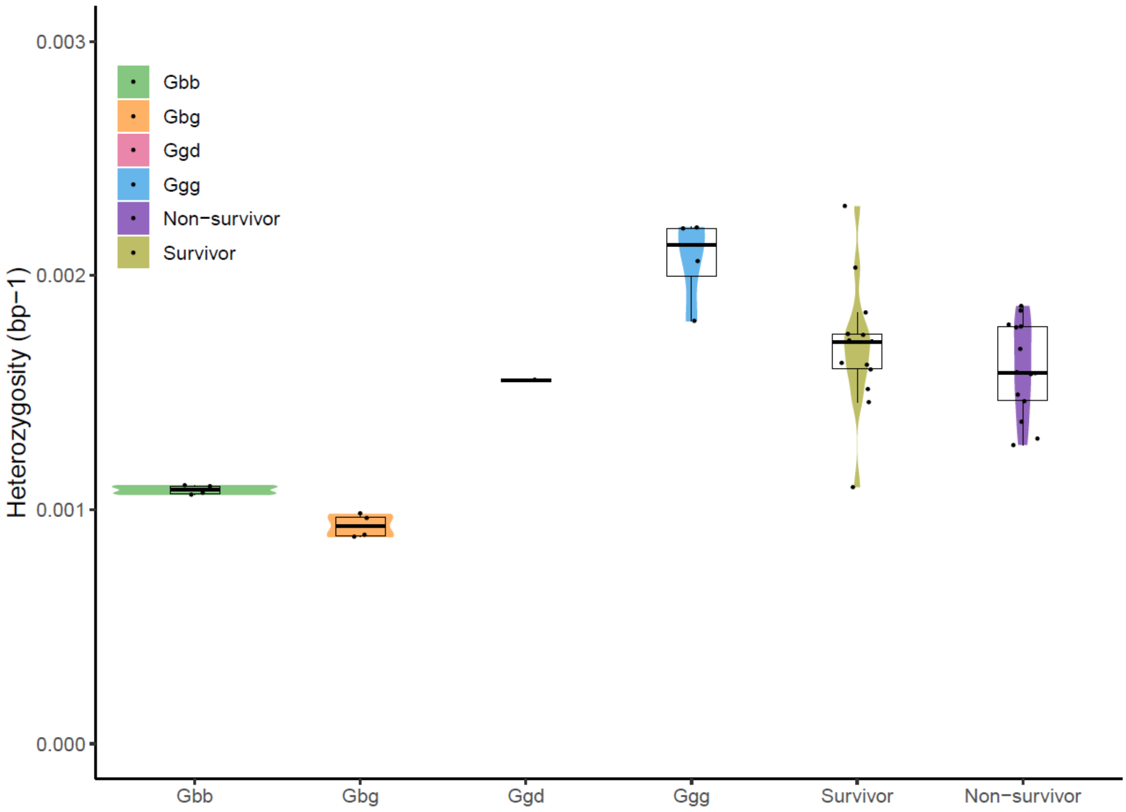


Fig. S6. Global heterozygosity for the target region of previously published gorilla genomes, together with the survivors and non-survivors of the Ebola outbreak. Gbb: *Gorilla beringei beringei*; Gbg: *Gorilla beringei graueri*; Ggd: *Gorilla gorilla dielhi*; Ggg: *Gorilla gorilla gorilla*.

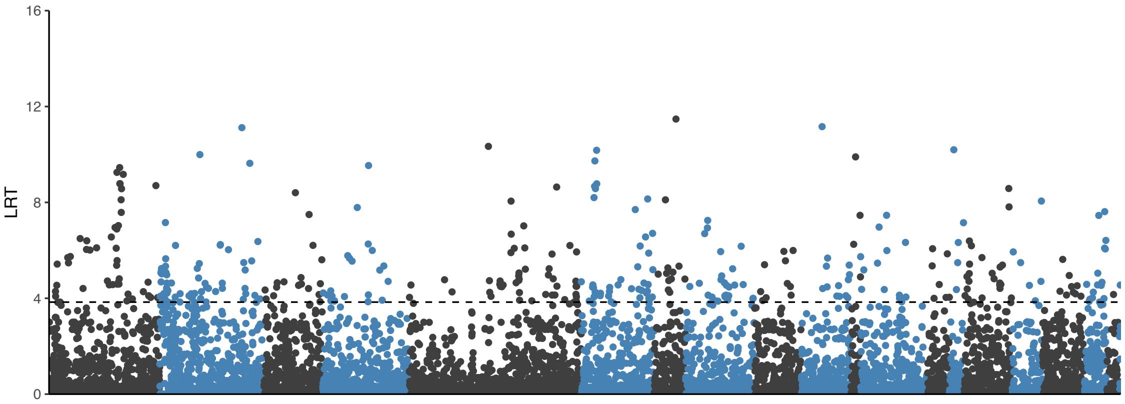


Fig. S7. Association analysis from ANGSD software. LRT value is the likelihood ration statistics which is chi square distributed with one degree of freedom. The significance threshold is set at 95% confidence, and thus the cut-off value is 3.84.

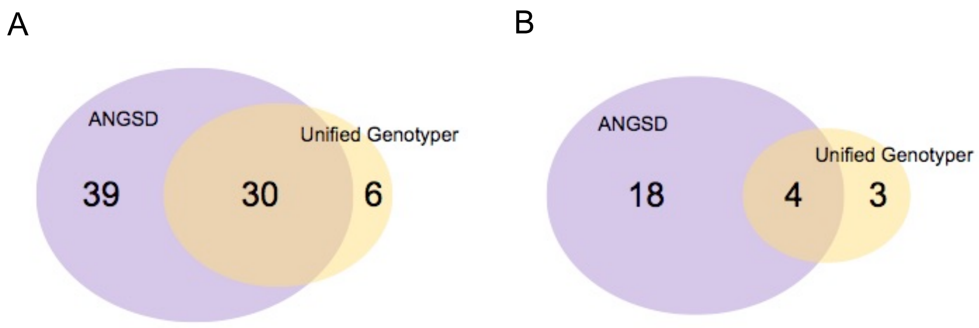


Fig. S8. Intersection of discovered genes with Unified Genotyper and with ANGSD. A) All discovered genes considered regardless of how many SNPs are significant in each gene. The intersection of both approaches is the following list of genes: *GBP1, CD58, CD1B, IGKJ4, CXCL2, NFKB1, IL2, BASP1, HLA-A, HLA-E, HLA-DRB5, ACTB, RAC1, IL6, ZNF398, LYN, TEK, ADAMTS13, IFIT2, ATM, DCTN2, BTG1, IGHJ3, B2M, ISG20, PFN1, CD68, CCL8, ERBB2* and *MX1*. B) Only discovered genes with at least 3 nominally significant SNPs. The intersection of both approaches is the following list of genes: *CD1B, ACTB, LYN* and *MX1*.

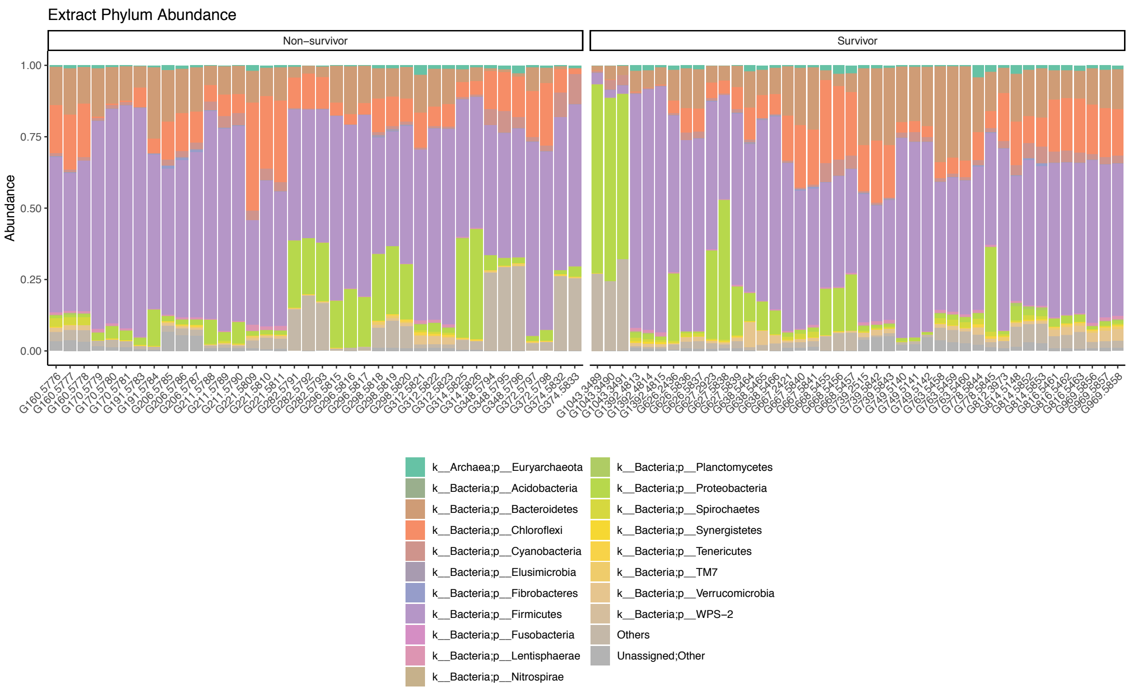


Fig. S9. Relative 16S rRNA taxa abundance (Phylum) for each fecal DNA extractions of the studied gorillas.

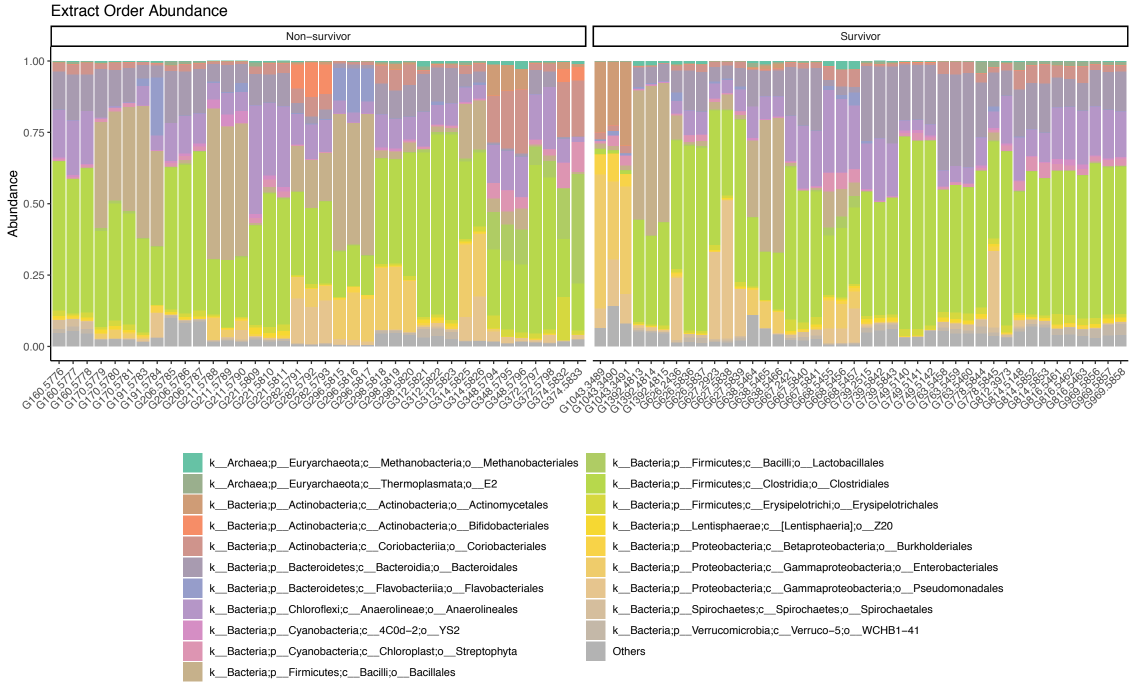


Fig. S10. Relative 16S rRNA taxa abundance (Order) for each fecal DNA extractions of the studied gorillas.

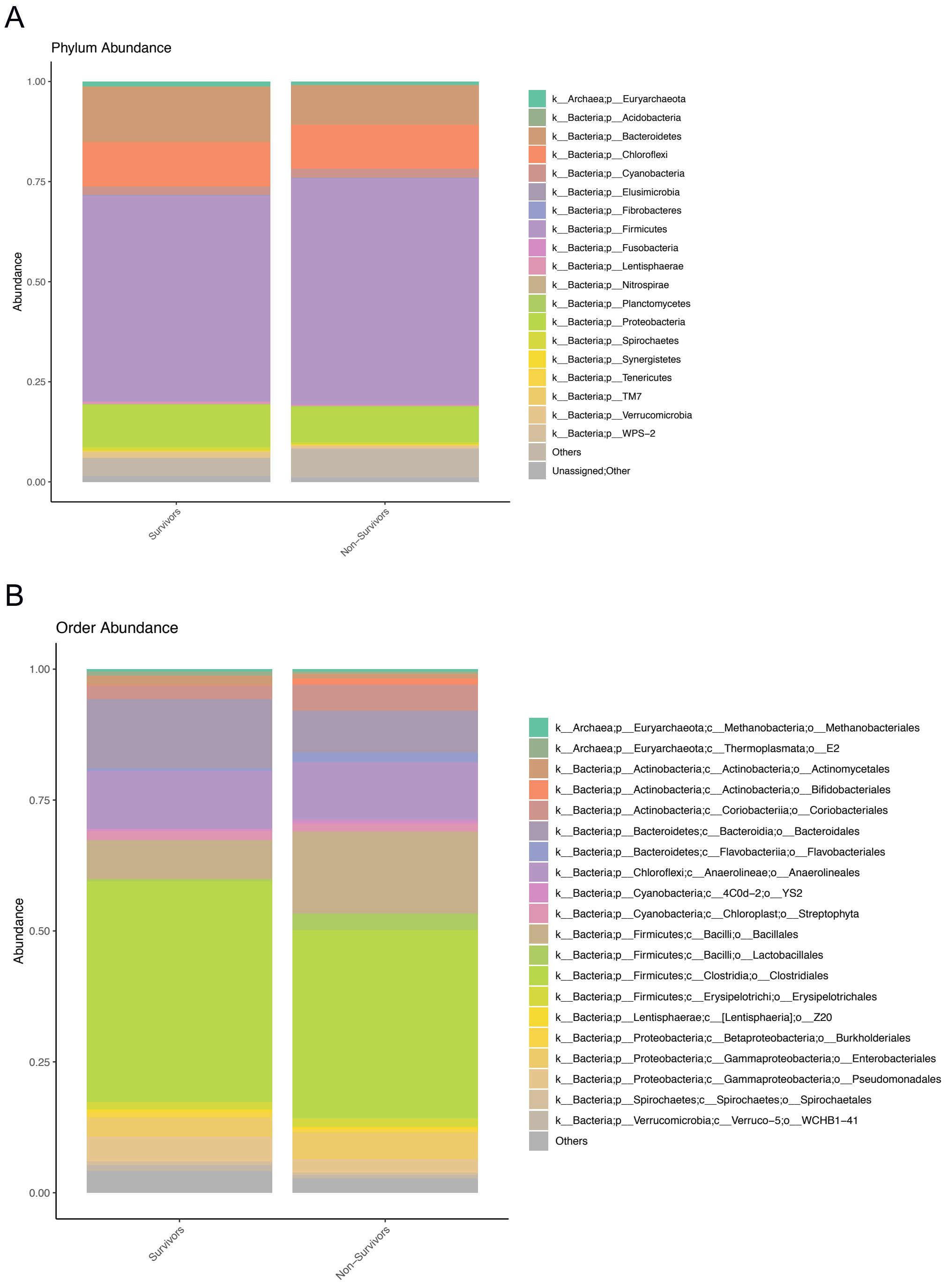


Fig. S11. Relative taxa abundance from the 16S rRNA microbiome dataset between Ebola outbreak gorilla survivors and non-survivors in A) phylum taxa and B) order taxa.

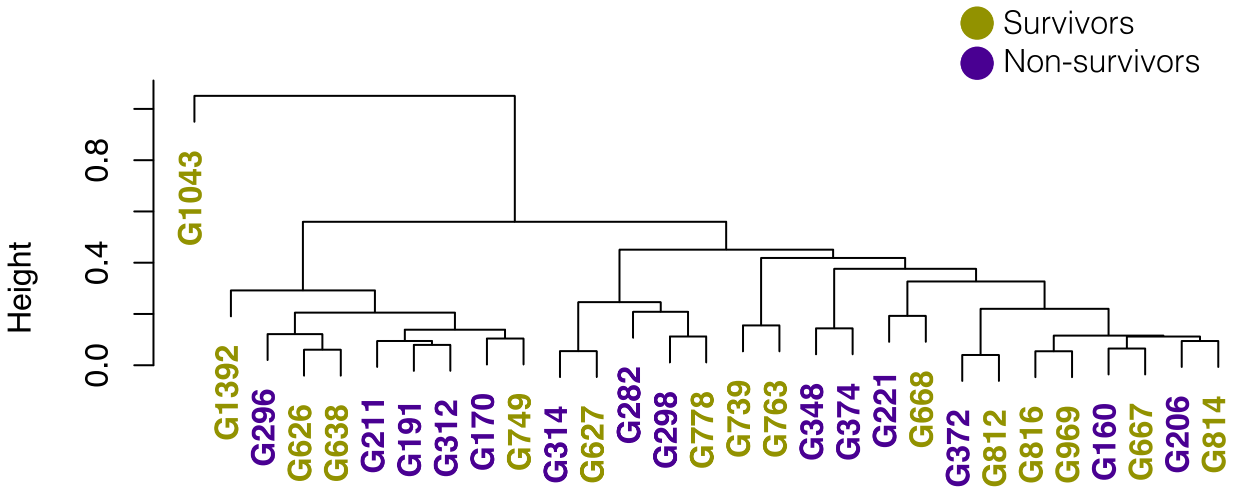


Fig. S12. Clustering dendrogram of the studied gorillas from the 16S rRNA microbiota analysis (order level) for all extractions of each sample.

A

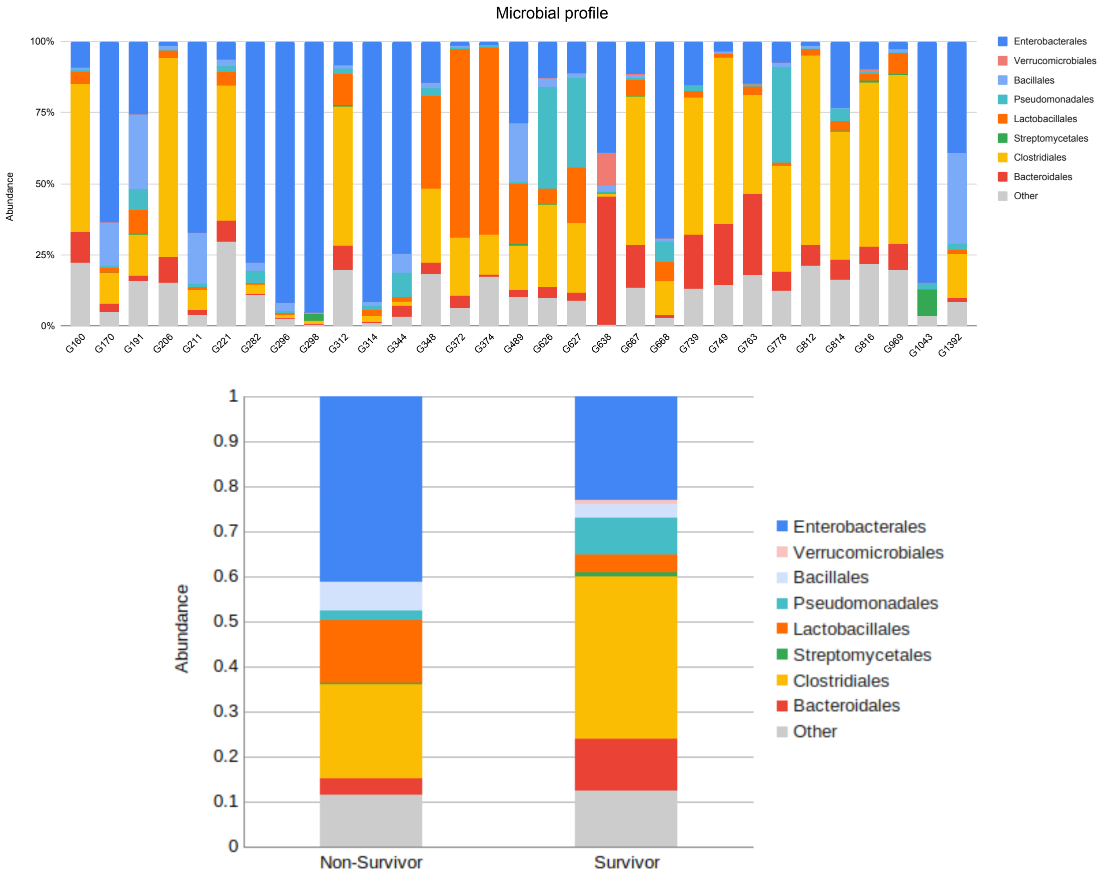


Fig. S13. Order level microbial profile of samples derived from shotgun microbial profiling. A) Per sample and B) grouping Survivors and Non-survivor gorilla individuals.

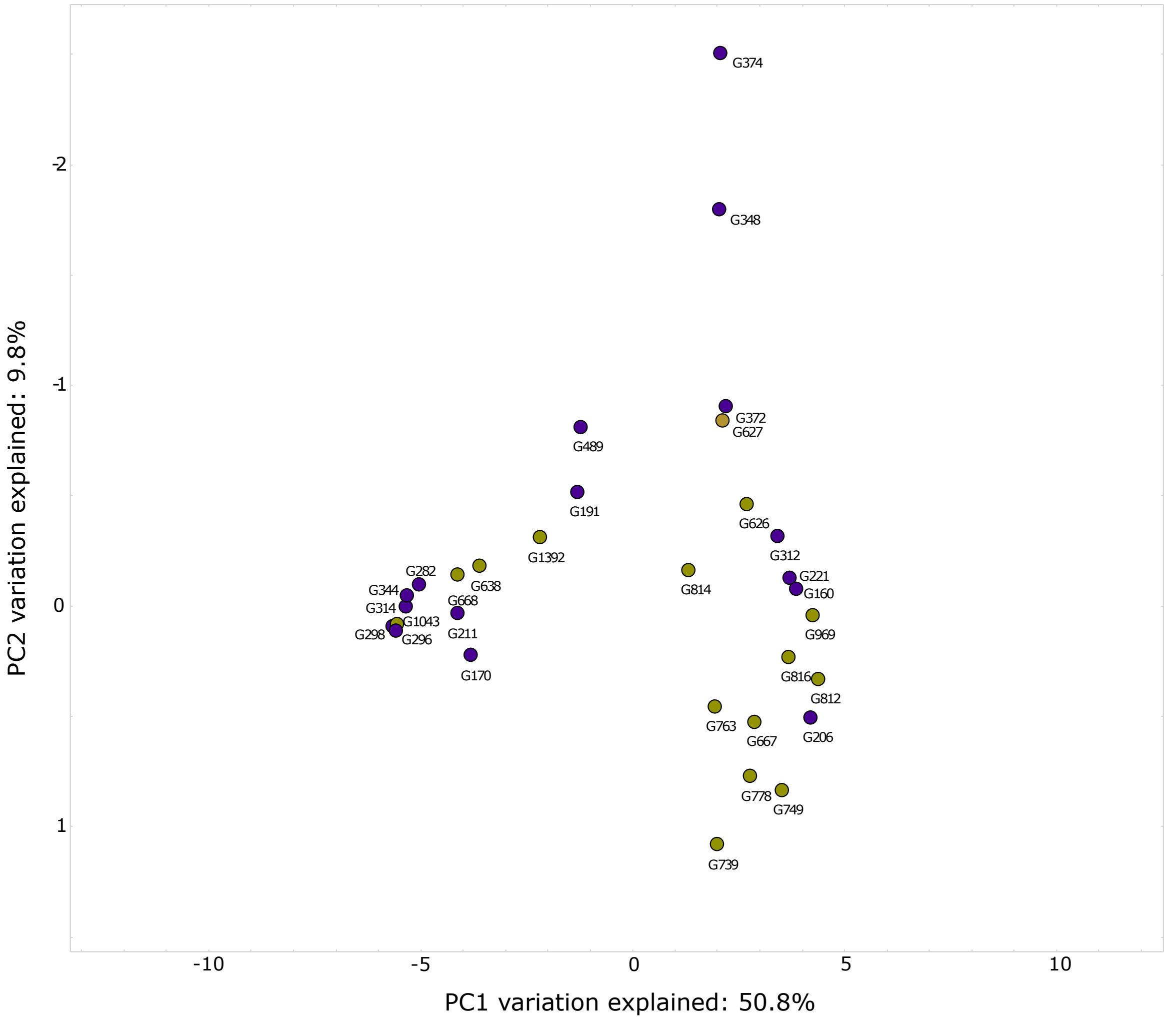


Fig. S14. Principal coordinate analysis of samples based on species composition, derived from shotgun microbial profiling. Ebola outbreak gorilla survivors are in green and the non-survivors in purple.

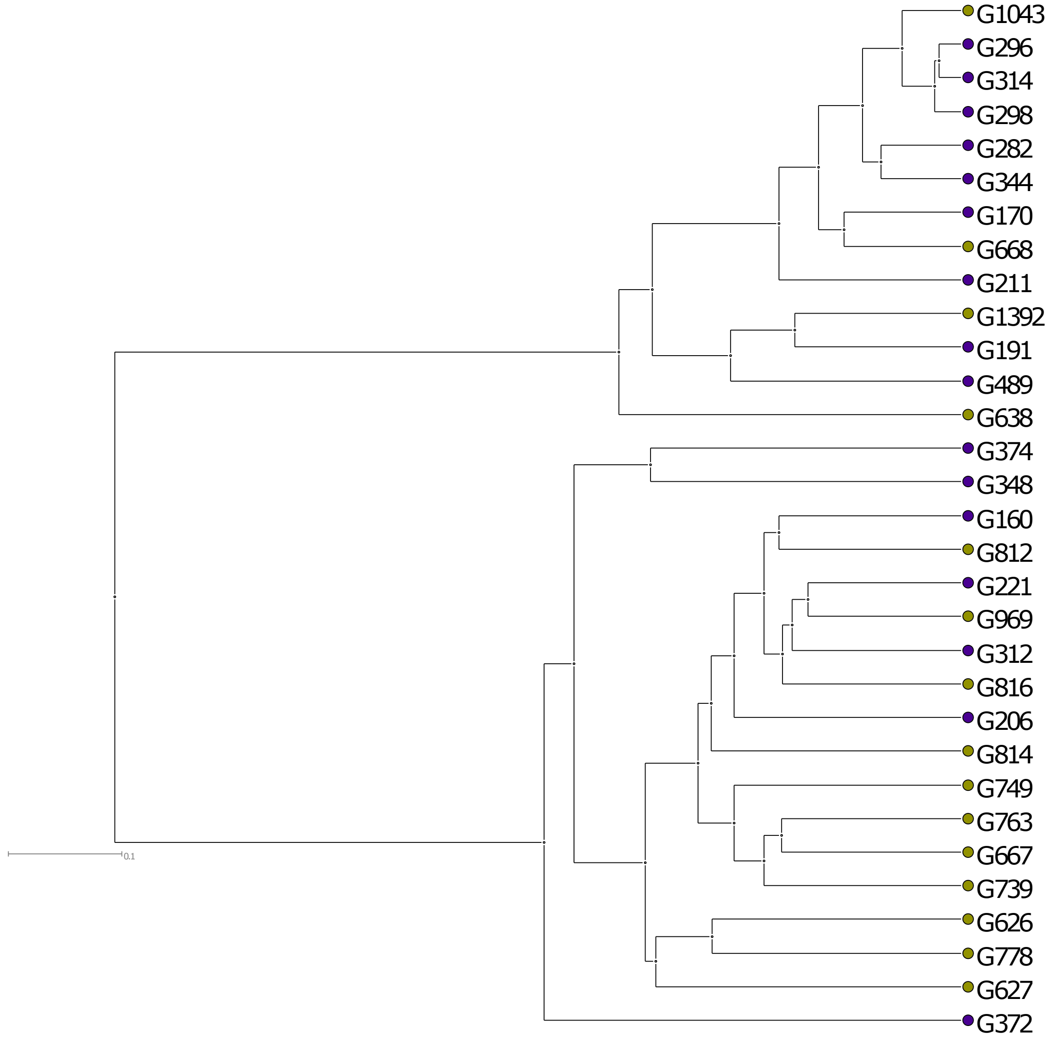


Fig. S15. Neighbor-joining tree of samples based on species composition, derived from shotgun microbial profiling. Ebola outbreak gorilla survivors are in green and the non-survivors in purple.
